# Molecular Screening of Hemoglobin S Variant in Anemia Patients of Eastern UP Population

**DOI:** 10.1101/830117

**Authors:** Vandana Rai, Upendra Yadav, Pradeep Kumar

## Abstract

Hemoglobinopathies are the most common type of inherited disease in human. in India the most frequent and clinically significant hemoglobin structural variants are HbS, HbD and HbE. The HbS mutation, in which a glutamic acid at position 6 in the β chain is substituted for valine Sickle cell disease is a major health problem in some parts of India. 2 ml blood sample was collected from 350 anemia patient and PCR-RFLP method was used for hemoglobin S analysis. Out of 350 samples, in four individuals, HbS mutation was found in homozygous (β ^6^/β ^6^) condition. All four individuals are Sickle cell cases. In conclusion, the percentage of Sickle cell disease was observed as 1.14% in Eastern UP anemic patients.

## Introduction

Abnormalities of hemoglobin (hemoglobinopathies) are the most common type of inherited disease in human. These variants are usually the result of point mutations in globin genes, commonly in β gene. However, in India the most frequent and clinically significant hemoglobin structural variants are HbS, HbD and HbE. The HbS mutation, in which a glutamic acid at position 6 in the β chain is substituted for valine (Codon 6 A-->T; β ^6^ Glu→Val), is responsible for sickle cell disease. Sickle cell disease affects the oxygen-carrying capacity of red blood cells. Valine is a hydrophobic amino acid, while glutamic acid is a hydrophilic amino acid. So this substitution creates a hydrophobic spot on the outside of the protein structure that sticks to the hydrophobic region of an adjacent hemoglobin molecule’s beta chain. This clumping together or polymerization of HbS molecules into rigid fibres causes the ‘sickling’ of red blood corpuscles (RBCs).

The highest frequency of HbS allele has been found 22.2% in Orissa followed by Madhya Pradesh (20%) Tamilnadu (20%), Assam (19.3%), Andhra Pradesh (18.3%) and Uttar Pradesh (9%) (Balgir, 1996). Now it is apparent that the sickle cell gene is not confined only to certain pockets of central and south India as previously postulated (Saha and Baneerji, 1973; Sharma, 1983), but it is prevalent throughout India, in almost all the states of India. In addition, the sickle cell gene is not only prevalent in the tribal or scheduled caste population of India (Saha and Baneerji, 1973; Sharma, 1983) but it has penetrated even into the general caste of India (Balgir and Sharma, 1988).

## Methods

The present study was approved by the Institutional Ethics Committee of VBS Purvanchal University, Jaunpur, India and all participants gave their written informed consent. 2 ml blood sample was collected from 350 anemia patients. Out of 350 samples, 50 samples were of Muslims, 100 from Scheduled cast, 100 from Brahmin and 100 samples from OBC. Genomic DNA was extracted using the standard method of Bartlett and White (2003). PCR was carried out using codon 6 specific PCR and amplicon was subsequently digested by DdeI restriction enzymes.

## Results and Discussion

HbS specific primers amplified 443 long DNA amplicon (Figure 1). In case of normal β-globin normal allele, DdeI enzyme cleave 443 bp amplicon in to two 376 and 67bp long fragments. HbS mutation creates an additional site and DdeI digestion of mutant allele produces three fragments of 201, 175 and 67 bp long fragments (Figure 2). Out of 350 samples, in four individuals, HbS mutation was found in homozygous (β ^6^/β ^6^) condition. All four individuals are Sickle cell cases. In 350 analyzed blood samples, heterozygous i.e sickle ell trait was not found in any sample. The overall HbS disease frequency was observed as 1.14% in Eastern UP (Figure 2). Out of four sickle cell cases, 1 individual was from Muslim religion, 2 individuals were SC, and 1 individual was Brahmin. The β^S^ frequency in Muslim, SC, and Brahmin caste were observed as 1%, 2%,and 1% respectively. The results of present study were well comparable with the findings of previously published article and it was also concluded that the β^S^ allele is also penetrated in general caste group (Balgir, 1996; Balgir and Sharma,1988).

**Figure 1.**
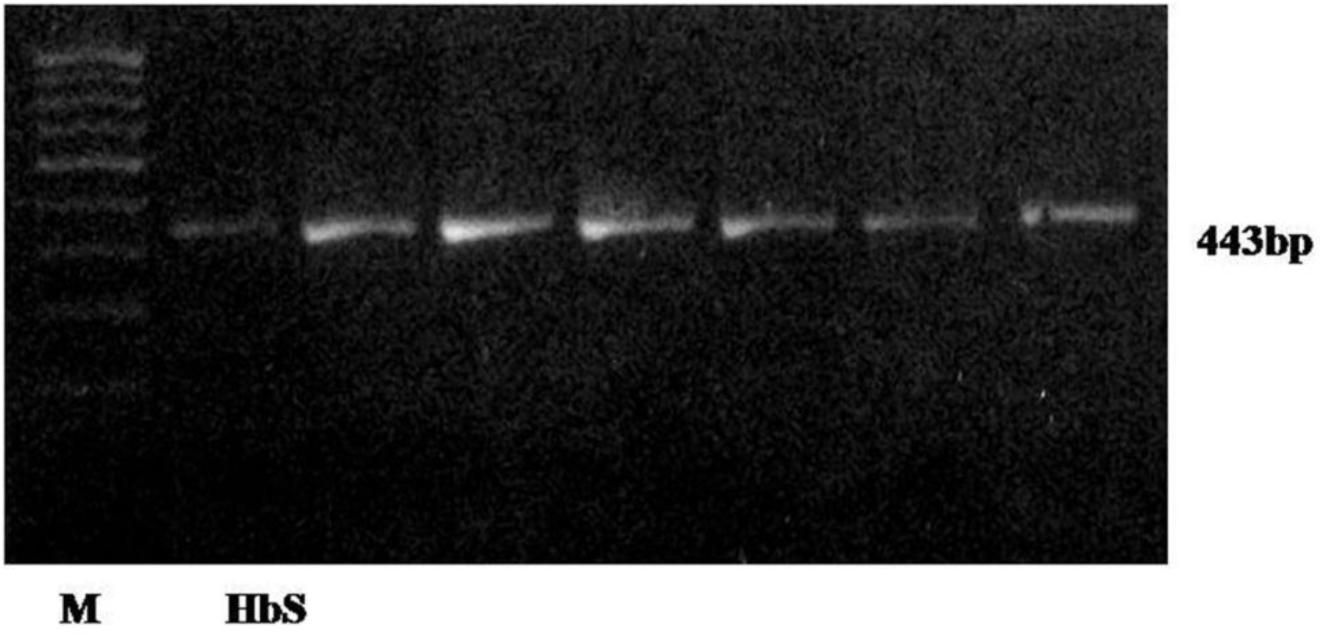
Gel picture showing 443bp long HbS amplicon.

**Figure 2.**
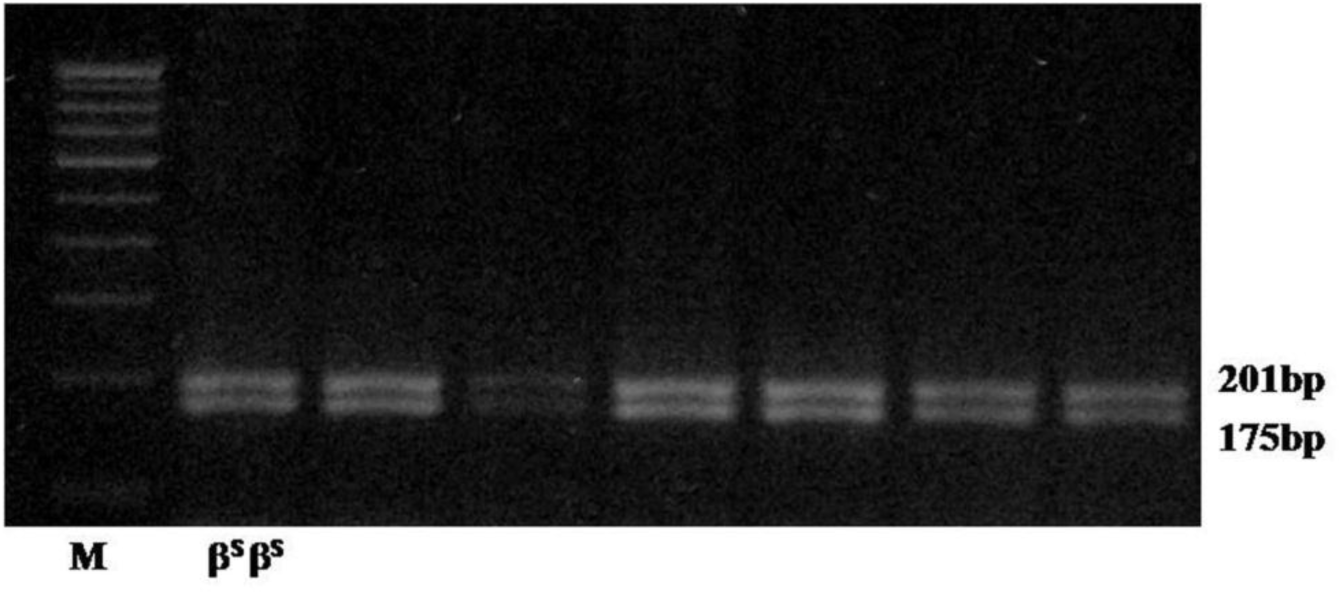
Gel picture showing DdeI digested amplicon.

Sickle cell disease is a major health problem in some parts of India that has great impact on both individuals and society. Newborn blood screening has been already adopted in the several countries to control the common inherited disease. It is possible to reduce morbidity and mortality rates in the first five years of life through neonatal screening program.

## Acknowledgements

We wish to thank all of the patients who participated in this study. This work was supported by a grant from Council of Scientific and Industrial Research, Government of India to Vandana Rai (No 27(0204)/09/EMR-II).

